# Characterization of fluorescent proteins, promoters, and selectable markers for applications in the Lyme disease spirochete *Borrelia burgdorferi*

**DOI:** 10.1101/363390

**Authors:** Constantin N. Takacs, Zachary A. Kloos, Molly Scott, Patricia A. Rosa, Christine Jacobs-Wagner

**Affiliations:** Microbial Sciences Institute, Yale West Campus, West Haven, CT, 06516, USA.; Department of Molecular, Cellular, and Developmental Biology, Yale University, New Haven, CT, 06511, USA.; Howard Hughes Medical Institute, Yale West Campus, West Haven, CT, 06516, USA.; Microbiology Program, Yale University, New Haven, CT, 06511, USA.; Laboratory of Bacteriology, Rocky Mountain Laboratories, Division of Intramural Research, National Institute of Allergy and Infectious Diseases, National Institutes of Health, Hamilton, MT, 59840, USA; Department of Microbial Pathogenesis, Yale School of Medicine, New Haven, CT, 06536, USA

## Abstract

Lyme disease is the most widely reported vector-borne disease in the United States. Its incidence is rapidly increasing and disease symptoms can be debilitating. The need to understand the biology of the disease agent, the spirochete *Borrelia burgdorferi*, is thus evermore pressing. Despite important advances in *B. burgdorferi* genetics, the array of molecular tools available for use in this organism remains limited, especially for cell biological studies. Here, we adapt a palette of bright and mostly monomeric fluorescent proteins for versatile use and multi-color imaging in *B. burgdorferi*. We also characterize two novel antibiotic selection markers and establish the feasibility of their use in conjunction with extant markers. Lastly, we describe a set of constitutively active promoters of low and intermediate strengths that allow fine-tuning of gene expression levels. These molecular tools complement and expand current experimental capabilities in *B. burgdorferi*, which will facilitate future investigation of this important human pathogen. To showcase the usefulness of these reagents, we used them to investigate the subcellular localization of BB0323, a *B. burgdorferi* lipoprotein essential for survival in the host and vector environments. We show that BB0323 accumulates at the cell poles and future division sites of *B. burgdorferi* cells, highlighting the complex subcellular organization of this spirochete.

**IMPORTANCE:** Genetic manipulation of the Lyme disease spirochete *B. burgdorferi* remains cumbersome, despite significant progress in the field. The scarcity of molecular reagents available for use in this pathogen has slowed research efforts to study its unusual biology. Of interest, *B. burgdorferi* displays complex cellular organization features that have yet to be understood. These include an unusual morphology and a highly fragmented genome, both of which are likely to play important roles in the bacterium’s transmission, infectivity, and persistence. Here, we complement and expand the array of molecular tools available for use in *B. burgdorferi* by generating and characterizing multiple fluorescent proteins, antibiotic selection markers, and constitutively active promoters of different strengths. These tools will facilitate investigations in this important human pathogen, as exemplified by the polar and midcell localization of the cell envelope regulator BB0323, which we uncovered using these reagents.

## INTRODUCTION

Lyme disease, a widespread infection transmitted by hard ticks of the *Ixodes* genus, is the most prevalent vector-borne disease in the United States. The disease is also common in Europe and Asia, and its incidence and geographic distribution have been steadily increasing in recent decades (1). Lyme disease is caused by spirochetal bacteria belonging to the *Borrelia burgdorferi* sensu lato group, with *B. burgdorferi* sensu stricto (hereafter referred to as *B. burgdorferi*) being the principal agent in North America, and *B. afzelii* and *B. garinii* being the primary agents in Eurasia. In humans, acute Lyme disease is often associated with a characteristic skin rash and flu-like symptoms. If left untreated, late stages of infection may result in carditis, neurological manifestations, and arthritis (2).

Spirochetes in general, and the *Borrelia* species in particular, display cellular features unusual for bacteria (3). Spirochete cells are typically very long and thin by bacterial standards. *B. burgdorferi* cells, for example, are 10 to 25 μm long and ~250 nm wide (4-6). Spirochetes are also highly motile, but, unlike most bacteria, their flagella are not external organelles (7). Instead, these flagella are located in the periplasm (i.e., between the inner and outer membranes). In *B. burgdorferi*, the helicity of the flagella imparts the flat-wave morphology of the bacterium (8). *B. burgdorferi* also possesses what is likely the most segmented genome of any bacterium investigated to date. It is made up of a linear chromosome of about 900 kilobases (kb) and over 20 linear and circular genetic elements ranging from 5 to 60 kb in length (9, 10). These smaller genetic elements are often referred to as plasmids, though many of them encode proteins that are essential for the life cycle of this organism (11). Recent work from our laboratory has shown that *Borreliae* species also have an uncommon pattern of cell wall synthesis in which discrete zones of cell elongation in one generation predetermine the division sites of daughter cells in the next generation (6).

While these unusual cellular features are integral to *B. burgdorferi* physiology and pathogenesis, little is known about how they arise or are maintained over generations. In fact, the cell biology of this pathogen remains largely unexplored. Technical hurdles have slowed progress in this area. Genetic manipulation of *B. burgdorferi* is feasible, but the available genetic tools are still limited, and the process remains cumbersome (12, 13). Constitutive gene expression is mostly limited to the use of very strong promoters. Moreover, apart from a few exceptions (14-19), fluorescent protein reporters have primarily been used as gene expression reporters or as cellular stains for *in vivo* localization of the spirochete (13). Yet fluorescent proteins have many more uses, which have transformed the field of cell biology (20). For example, fluorescent proteins have opened the door to localization studies in live cells. They have also facilitated the detection of protein-protein interactions, the measurement of physical properties of cells, and the investigation of single events and of population heterogeneity. Much of this information is not accessible through the use of bulk biochemical measurements on cell populations. The averaging inherent to such techniques leads to loss of spatial resolution and obscures rare events and cell-to-cell or subcellular heterogeneity of behaviour (21). Indeed, the ability to perform extensive genetic manipulations and to use a wide panel of fluorescent proteins in an organism has been key to progress in understanding bacterial cell biology (22). Such approaches have been used extensively in model bacteria such as *Bacillus subtilis, Escherichia coli*, and *Caulobacter crescentus* since the first reported use of fluorescent protein fusions two decades ago (23-25). In order to facilitate the study of *B. burgdorferi*, we have generated new investigative tools by characterizing a panel of fluorescent proteins, promoters and antibiotic resistance markers for use in this medically important bacterium. We exemplify the usefulness of these reagents by creating an mCherry fusion to BB0323, a multifunctional *B. burgdorferi* lipoprotein required for outer membrane stability (26-28) and essential for the spirochete’s survival in the tick vector and the mammalian host (27). Using this fusion, we show that BB0323 localizes at the spirochete’s poles and at future division sites, highlighting underappreciated spatial and temporal organization principles of *B. burgdorferi* cells. (An earlier version of this article was submitted to the online preprint archive BioRxiv [doi: https://doi.org/10.1101/363390])

## MATERIALS AND METHODS

### Bacteria, growth conditions, and genetic transformations

Bacterial strains used in this study are listed in Table 1. *E. coli* strains were grown at 30 °C in liquid culture in Super Broth medium (35 g/L bacto-tryptone, 20 g/L yeast extract, 5 g/L NaCl, 6 mM NaOH) with shaking, or on LB agar plates. Plasmids were transformed by electroporation or heat shock. For selection of *E. coli* strains we used 200 μg/mL (solid medium) or 100 μg/mL (liquid medium) ampicillin, 20 μg/mL (solid medium) or 15 μg/mL (liquid medium) gentamicin, 50 μg/mL kanamycin (solid and liquid media), 50 μg/mL spectinomycin (solid medium), 50 μg/mL streptomycin (liquid medium), and 25 μg/mL (liquid medium) or 50 μg/mL (solid medium) rifampicin.

**Table 1.**
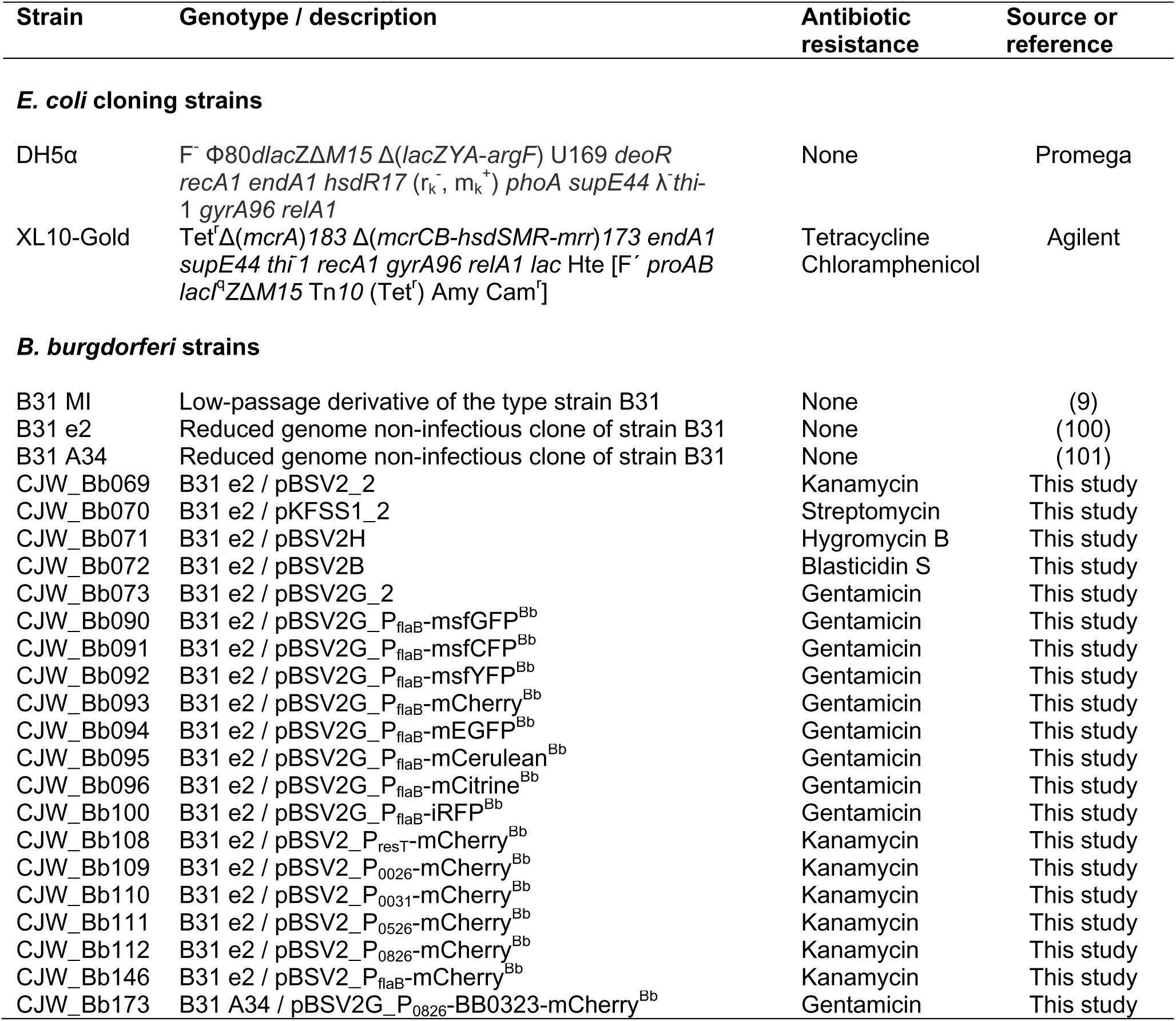
Strains used in this study

*B. burgdorferi* strains were grown in BSK-II medium supplemented with 6% (v/v) heat inactivated rabbit serum (Sigma Aldrich or Gibco) or in complete BSK-H medium (Sigma Aldrich), as previously described (29-31). Cultures were incubated at 34 °C under 5% CO_2_ atmosphere in a humidified incubator. Antibiotics were used at the following concentrations (unless otherwise indicated): gentamicin at 40 μg/mL, streptomycin at 100 μg/mL, kanamycin at 200 μg/mL, blasticidin S at 10 μg/mL, and hygromycin B at 250 μg/mL. Ampicillin was purchased from Fisher Scientific, blasticidin S and hygromycin B from Invivogen, and all other antibiotics and biliverdin hydrochloride from Sigma Aldrich.

### *B. burgdorferi* strain generation

*B. burgdorferi* electrocompetent cells were prepared as previously described (32) and were transformed with shuttle vector plasmid DNA (usually 30 μg) by electroporation. Electroporated cells were then allowed to recover overnight in BSK-II medium at 34 °C. The next day, the transformants were plated in semi-solid BSK-agarose medium with appropriate antibiotics as previously described (32). Individual colonies were then expanded and characterized. Alternatively, antibiotic selection was initiated in liquid medium and 5-fold serial dilutions of the culture were plated in a 96-well plate (24 wells for each dilution). After 10-14 days of incubation, the wells were inspected by microscopy using dark-field illumination. Based on Poisson distribution probability, when fewer than 20% of the wells of a given dilution were positive for growth, those wells were considered to contain clonal populations and were further expanded and characterized as such. When appropriate, fluorescence imaging was used to confirm fluorescent protein expression. Alternatively, selected, non-clonal transformant populations were enumerated using C-Chip disposable hemocytometers (INCYTO), using the manufacturer’s instructions with the following change: counting was done by continuously scanning the full height of the counting chamber for each counting surface to account for the height of the counting chamber being larger than the size of the spirochetes. Enumerated spirochetes were then diluted in BSK-II medium and plated in 96-well plates at an average density of 0.2 cells/well. After 10-14 days, clonal growth was confirmed by dark-field microscopy imaging.

### Determination of minimal inhibitory concentrations (MIC) and antibiotic cross-resistance

MICs were determined using strains B31 e2 or B31 MI, while cross-resistance testing was done using B31 e2-derived strains that contained shuttle vectors carrying kanamycin, gentamicin, streptomycin, blasticidin S, or hygromycin B resistance markers (see strains CJW_Bb069 through CJW_Bb073 in Table 1). For both tests, antibiotics were two-fold serially diluted in complete BSK-II or BSK-H medium. For each concentration, 100 μL of antibiotic solution were dispensed into two to four wells of 96-well plates. The cell density of *B. burgdorferi* cultures was determined by direct counting using dark-field microscopy. The cultures were then diluted to 2 ×10^4^ cells/mL in antibiotic-free medium, and 100 μL of this diluted culture were added to the antibiotic-containing wells to yield an inoculum of 10^4^ cells/mL. The plates were incubated for at least 4 days at 34 °C under 5% CO_2_ atmosphere in a humidified incubator, after which each well was checked for spirochete growth and motility using dark-field microscopy. A well was marked as positive if motile cells were detected. The plates were further incubated for several days, during which bacterial growth-dependent acidification caused the phenol red pH indicator in the medium to change color. This color change was documented using colorimetric trans-illumination imaging on a GE Amersham Imager 600. We verified that growth scoring of each well by dark-field imaging matched the observed medium color change.

### DNA manipulations

Plasmids used in this study are listed in Table 2. Methods of plasmid construction and sequences of oligonucleotide primers are provided as Supplemental Material. Standard molecular biology techniques were used, as detailed in the Supplemental Material. Codon optimization was performed using the web-based Java Codon Adaptation Tool hosted at www.jcat.de (33) and the codon usage table for *B. burgdorferi* as stored at www.kazusa.or.jp/codon (34). Codon-optimized DNA sequences were then chemically synthesized at Genewiz. DNA sequences of each codon-optimized gene are provided in the Supplemental Material. They are also available at GenBank under the accession numbers MH644044 through MH644053. The names of these genes include a *Bb* superscript to indicate that the gene’s nucleotide sequence is codon-optimized for translation in *B. burgdorferi* (e.g., *iRFP^Bb^*). The name of the protein encoded by such a gene (e.g., iRFP), however, does not include the Bb superscript, as the protein’s amino acid sequence does not differ from that expressed from other versions of the gene.

**Table 2.**
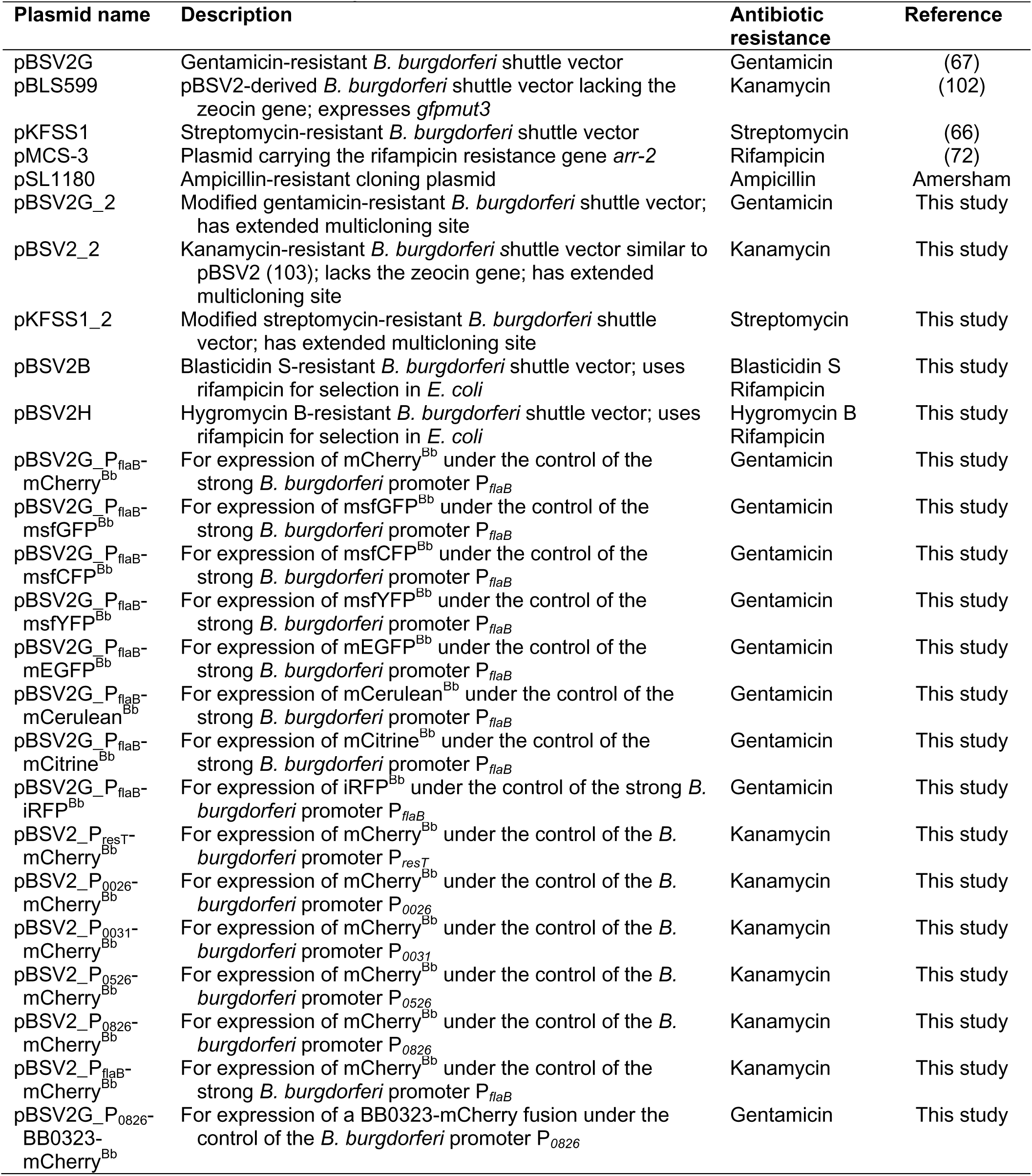
Plasmids used in this study.

### Microscopy

Visualization and counting of live spirochetes were done using a Nikon Eclipse E600 microscope equipped with dark-field illumination optics and a Nikon 40X 0.55 numerical aperture (NA) phase contrast air objective. Phase contrast and fluorescence imaging was done on a Nikon Eclipse Ti microscope equipped with a 100X Plan Apo 1.40 NA phase contrast oil objective, a Hamamatsu Orca-Flash4.0 V2 Digital CMOS camera, a Sola light engine (Lumencor), and controlled by the Metamorph software (Molecular Devices). Alternatively, light microscopy was performed on a Nikon Ti microscope equipped with a 100X Plan Apo 1.45 NA phase contrast oil objective, a Hamamatsu Orca-Flash4.0 V2 CMOS camera, a Spectra X light engine (Lumencor) and controlled by the Nikon Elements software. Excitation of iRFP was achieved using the 640/30 nm band of the SpectraX system, but higher excitation efficiency and thus brightness could in theory be obtained using a red-shifted excitation source between 660 and 680 nm. The following Chroma filter sets were used to acquire fluorescence images: CFP (excitation ET436/20x, dichroic T455lp, emission ET480/40m), GFP (excitation ET470/40x, dichroic T495lpxr, emission ET525/50m), YFP (excitation ET500/20x, dichroic T515lp, emission ET535/30m), mCherry/TexasRed (excitation ET560/40x, dichroic T585lpxr, emission ET630/75m), and Cy5.5 (excitation ET650/45x, dichroic T685lpxr, emission ET720/60m). For imaging, cultures were inoculated at densities between 10^3^ and 10^5^ cells/mL, and grown for two to three days to reach densities between 10^6^ and 3 × 10^7^ cells/mL. The cells were then immobilized on a 2% agarose pad (6, 35) made with phosphate buffered saline covered with a No. 1.5 coverslip, after which the cells were immediately imaged live. Images were processed using the Metamorph software. Figures were generated using Adobe Illustrator software.

### Image analysis

Cell outlines were generated using phase contrast images and the open-source image analysis software Oufti (36). Outlines were checked visually for each cell and were extended manually to the full length of the cells when appropriate. When not assigned to single cells or assigned to non-cellular debris, outlines were manually removed. The remaining outlines were further refined using the Refine All function of the software. To quantify fluorescence signals, individual cytoplasmic cylinders connected by an outer membrane bridge (i.e., late predivisional cells with two separated cytoplasms) were treated as independent cellular units. For demograph analysis of strain CJW_Bb173, late predivisional cells were considered to form one cell. Fluorescence signal data was added to the cells and demographs were generated in Oufti. The resulting cell lists were processed using the MATLAB script addMeshtoCellList.m (see Supplemental Material for the code). This script uses the functions CL_getframe.m, CL_removeCell.m, CL_cellId2PositionInFrame.m, and getextradata.m, which were previously described (36). Single-cell fluorescence intensity values were calculated by dividing the total fluorescence signal inside a cell outline by the cell’s area using the MATLAB-based function CalculateFluorPerCell.m. Final fluorescence data were plotted using the GraphPad Prism 5 software. The number of cells analyzed for each condition is provided in the Supplemental Material.

## RESULTS

### A wide palette of fluorescent proteins for imaging in *B. burgdorferi*

Only a few fluorescent proteins have been used to date in *B. burgdorferi* (summarized in Table 3). These proteins belong primarily to two color classes: green fluorescent proteins (GFP) and red fluorescent proteins (RFP) (Table 3). To expand the range of options for multi-color imaging of *B. burgdorferi*, we focused on a set of fluorescent proteins that have been used in localization studies in other organisms and codon-optimized their genes for translation in *B. burgdorferi*. The selected proteins span five color classes (Table 3) and their signal can be collected using widely available filter sets for cyan fluorescent protein (CFP), GFP, yellow fluorescent protein (YFP), mCherry/TexasRed and Cy5.5 fluorescence. The selected cyan, green, and yellow variants are all derivatives of the jellyfish (*Aequorea victoria*) GFP. We used both the classic variants Cerulean (37), enhanced GFP, or EGFP (38), Citrine (39), as well as the superfolder (e.g., sfGFP) variants (40). All variants included the monomeric mutation A206K (41), denoted by a lower case m before the name of the protein (e.g., mCerulean). Our red protein of choice was mCherry (42), a monomeric, improved variant of mRFP1. Lastly, we codon-optimized and expressed an infrared fluorescent protein, iRFP (43). The far-red wavelengths used to excite this fluorophore are less toxic to cells than the shorter excitation wavelengths used for the other fluorescent proteins, and the sample autofluorescence in the near-infrared spectral region is lower than in the other, blue-shifted imaging windows (20, 44).

**Table 3.** Fluorescent proteins used in *B. burgdorferi*

| Color class | Protein expressed | Ex/Em<sup>a</sup> max (nm) | Source or reference for protein/gene development | Reference for use in *B. burgdorferi* | Notes |
| --- | --- | --- | --- | --- | --- |
| **I. Fluorescent proteins previously used in *B. burgdorferi*:** |  |  |  |  |  |
| Cyan | CFP | 434/477<sup>b</sup> | Clontech; (104) | (105) | Rarely used |
| Green | EGFP | 489/509 | Clontech; (38) | (68) | Low expression; has mammalian codon usage |
|  | GFPmut1 | 488/507 | (104, 106-108) | (105) | Widely used; adapted for bacterial expression; same protein as EGFP |
|  | GFPmut3 | 501/511 | (106) | (102) |  |
|  | GFP cycle 3 | NR<sup>c</sup> | (109) | (110) | Retains UV excitation peak |
| Yellow | YFP | 514/527<sup>b</sup> | (104) | (105) | Rarely used |
| Red | mRFP1 | 584/607 | (111) | (18) | Folds in the periplasm |
|  | dTomato | 554/581 | (42) | (112) | Dimeric |
| **II. New fluorescent proteins adapted for use in *B. burgdorferi*:** |  |  |  |  |  |
| Cyan | mCerulean | 433/475 | (37) | This study | A206K monomeric mutation |
|  | msfCFP | NR<sup>c</sup> | (40) | This study | A206K mutation; superfolder |
| Green | mEGFP | 489/509 | (38) | This study | A206K mutation |
|  | msfGFP | 485/NR<sup>c</sup> | (40) | This study | A206K mutation; superfolder |
| Yellow | mCitrine | 516/529 | (39) | This study | A206K mutation |
|  | msfYFP | NR<sup>c</sup> | (40) | This study | A206K mutation; superfolder |
| Red | mCherry | 587/610 | (42) | This study |  |
| Infrared | iRFP | 690/713 | (43) | This study | Dimeric |

<sup>a</sup> Maximum excitation (Ex) and emission (Em) wavelengths; <sup>b</sup> values assumed to be those for ECFP and EYFP, respectively (20); <sup>c</sup>NR, not reported: values were not reported in the original publication, or could not be exactly inferred from excitation and emission graphs.

To visualize these fluorescent proteins, we expressed them in strain B31 e2 from the strong flagellin promoter P*_flaB_* (45) located on a shuttle vector. With the exception of iRFP, each fluorescent protein displayed bright fluorescence when imaged using a filter set matched to its color (Figure 1A). Unlike the other fluorescent proteins, which oxidatively conjugate their own amino acid side chains to create a fluorophore (20), iRFP covalently binds an exogenous biliverdin molecule, which then serves as the fluorophore (43). Adding the biliverdin cofactor to the growth medium of the iRFP-expressing strain rendered the cells fluorescent in the near-infrared region of the spectrum, as detected with a Cy5.5 filter set (Figure 1B). Treating a control strain carrying an empty shuttle vector with biliverdin did not cause any increase in cellular fluorescence (data not shown). To measure cellular fluorescence levels, we chose a microscopy-based approach in conjunction with quantitative image analysis. This allowed us to efficiently analyze hundreds of cells and to clearly distinguish individual cells from similarly-sized debris found in the culture medium, or from clumps of multiple cells. Using this method, we established that a 4 μM concentration of biliverdin in the growth medium was sufficient to achieve maximal cellular brightness (Figure 1C). Close-to-maximal iRFP brightness was reached as early as an hour after addition of biliverdin to the culture and was maintained throughout subsequent growth (Figure 1D). Furthermore, continuous growth of *B. burgdorferi* in the presence of biliverdin was indistinguishable from growth in biliverdin-free medium (Figure 1E). This indicates that culture experiments that involve iRFP may be performed either by adding biliverdin shortly before imaging or by growing the cells continuously in the presence of biliverdin.

**Figure 1.**
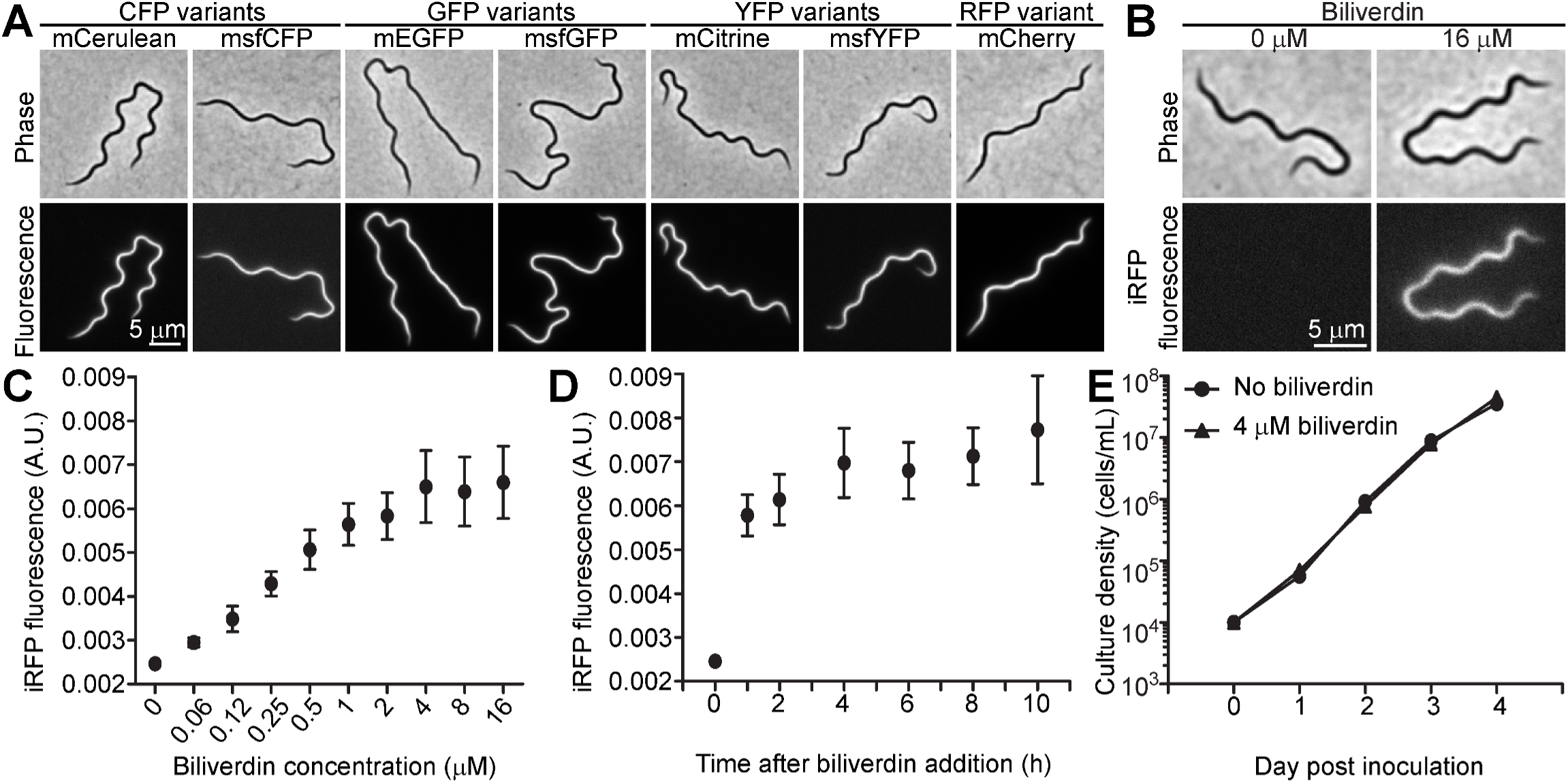
Fluorescent protein characterization. **A.** *B. burgdorferi* strains CJW_Bb090 through CJW_Bb096 expressing the indicated fluorescent proteins were imaged with matching filter sets. **B.** Strain CJW_Bb100 expressing iRFP requires biliverdin for development of fluorescence. Cells were grown in liquid culture with biliverdin for two days prior to imaging using a Cy5.5 filter set. **C.** Dose-response of iRFP fluorescence to biliverdin concentration. Strain CJW_Bb100 was grown in the presence of biliverdin for two days prior to imaging. Between 86 and 206 cells were analyzed for each concentration. Total cellular fluorescence levels were normalized by the cell area. Shown are means ± standard deviations (SD). A.U., arbitrary units. **D.** Time-course of iRFP fluorescence development in strain CJW_Bb100 following addition of 16 μM biliverdin. Between 68 and 110 cells were analyzed for each time point. **E.** Biliverdin does not affect *B. burgdorferi* growth. Strain CJW_Bb100 was inoculated at 10^4^ cells/mL in duplicate in medium containing 4 μM biliverdin or no biliverdin, after which the spirochetes were enumerated daily.

**Table 4.** Summary of antibiotic resistance markers used in *B. burgdorferi*^a^

| Resistance gene | Antibiotic | MIC<sup>b</sup><br>(µg/mL) | Notes | References |
| --- | --- | --- | --- | --- |
| **I. Widely used resistance markers** |  |  |  |  |
| *aphI* | Kanamycin | <25 | Cross-resistance to neomycin, lividomycin, paromomycin, ribostamycin | (45, 67) |
| *aadA* | Streptomycin | 7<sup>c</sup> | Expected cross-resistance to spectinomycin | (66) |
| *aacC1* | Gentamicin | <15.6 |  | (67) |
| *ermC* | Erythromycin | 0.005 | Resistance level varies among strains; May pose safety risk | (68, 113, 114) |
| **II. Newly developed resistance markers** |  |  |  |  |
| *bsd*<sup>Bb</sup> | Blasticidin S | <5 | Neither marker provides cross-resistance to the selection antibiotics listed above. | This study |
| *hph*<sup>Bb</sup> | Hygromycin B | <200 |  | This study |

<sup>a</sup>For space considerations, this table does not contain a comprehensive list of antibiotic resistance markers developed for use in *B. burgdorferi*. For a detailed discussion of other markers, please see (13); <sup>b</sup>Minimal inhibitory concentration, determined in liquid culture; <sup>c</sup>Value is that of an inhibitory dose 50, or ID<sub>50</sub>.

In microscopy studies, simultaneous imaging of multiple fluorescent proteins requires that the signal generated by a given fluorescent protein does not overlap with the fluorescence channels used to collect the signal of another protein. To assess the viability of using our palette of fluorescent proteins for multi-color imaging in *B. burgdorferi*, we quantified the signal generated by each fluorescent protein when imaged with the commonly used CFP, GFP, YFP, mCherry, and Cy5.5 filter cubes (Figure 2). We found that each fluorescent protein generated a strong signal when imaged with a color-matched filter set (Figure 2). As expected, we detected a significant spectral overlap between CFP and GFP, as well as between GFP and YFP variants. Importantly, signal quantification showed that mCerulean or msfCFP can be imaged alongside mCitrine, mCherry, and iRFP, while mEGFP or msfGFP can be imaged alongside mCherry and iRFP, opening the door to combinatorial imaging of up to four proteins in the same *B. burgdorferi* cell.

**Figure 2.**
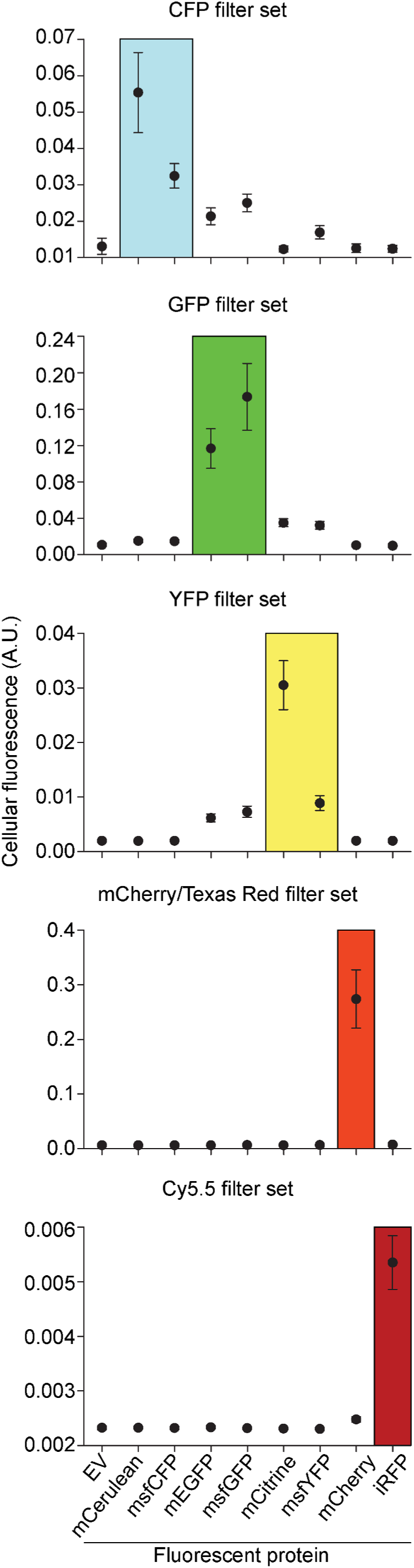
Quantification of fluorescent protein signal using common fluorescence filter sets. Strains CJW_Bb090 through CJW_Bb096 and CJW_Bb100 expressing the fluorescent proteins indicated at the bottom of the figure were each imaged using five filter sets: CFP, GFP, YFP, mCherry/TexasRed, and Cy5.5 (see the Materials and Methods section for filter set specifications). Strain CJW_Bb073 carrying an empty shuttle vector (EV) was also imaged to measure the cellular autofluorescence. Each filter set is listed at the top of the corresponding graph. Fluorescence intensity values were normalized by the cell area and are depicted as means ± SD in arbitrary units (A.U.). For each strain, 117 to 308 cells were analyzed. The iRFP strain was grown in the presence of 4 μM biliverdin for three days prior to imaging. The boxed region in each plot highlights the data obtained with filter sets that were ideal for the expressed fluorescent protein.

### Promoters for various levels of constitutive expression in *B. burgdorferi*

To date, constitutive expression of exogenous genes in *B. burgdorferi*, including antibiotic selection markers and reporter genes such as fluorescent proteins and luciferases, has almost exclusively relied on very strong promoters such as P*_flaB_* and P*_flgB_* (13, 45). Reporter expression from strong promoters facilitates spirochete detection, particularly in high fluorescence background environments such as the tick midgut or mammalian tissues (46-48). However, as overexpression can affect protein localization, interfere with function, or cause cellular toxicity (e.g., (49-59)), lower levels of gene expression have proven instrumental in facilitating localization studies (e.g., (60-63)) and are often preferred in such applications.

To identify constitutively expressed promoters of low and medium strengths, we mined a published RNA sequencing (RNA-seq) dataset that measured transcript levels in cultures of *B. burgdorferi* in early-exponential, mid-exponential and stationary phases of growth (64). We selected five genes whose expression was largely unchanged among the three growth phases tested (Figure 3A), amplified a DNA region upstream of each gene’s predicted translational start site, and fused it to an mCherry reporter in a kanamycin resistance-conferring shuttle vector (Figure 3B). The amplified putative promoter sequences ranged in size from 129 to 212 base pairs (bp) and included the reported 5’ untranslated regions (5’UTRs) of the downstream genes (64, 65). We also included in our analysis an empty vector and a vector containing a P*_flaB_*-*mCherry^Bb^* fusion, which served as references for no and high expression, respectively.

**Figure 3.**
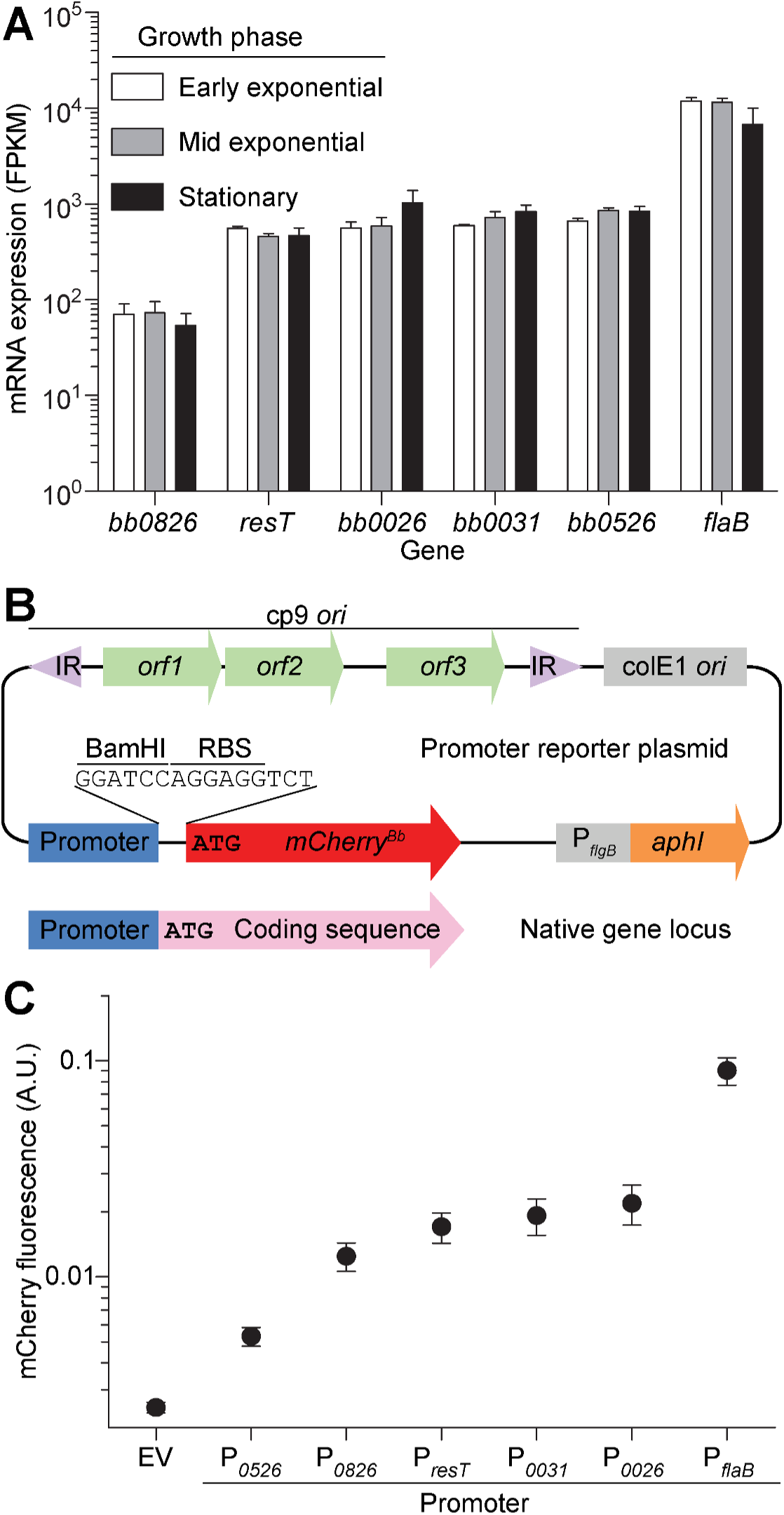
Promoter strength quantification. **A.** mRNA expression levels extracted from published RNA-seq data obtained using strain B31-A3 (99) grown to early exponential phase (10^6^ cells/mL), mid-exponential phase (10^7^ cells/mL), or stationary phase (one day after reaching 10^8^ cells/mL) (64). FPKM, fragments per kilobase transcript per million mapped reads. **B.** Promoter reporter plasmid map (not drawn to scale). IR, inverted repeats; cp9 *ori*, origin of replication of *B. burgdorferi* plasmid cp9, which includes the genes *orf1, orf2, orf3* needed for plasmid replication in *B. burgdorferi*; colE1 *ori, E. coli* origin of replication; P*_flgB_, B. burgdorferi* flagellar rod operon promoter; *aphI*, kanamycin resistance gene. The promoter (blue) and the mCherry-coding sequence (red) are connected by a BamHI restriction enzyme site and a ribosomal binding site (RBS). The native locus from which the promoter was extracted is depicted below the plasmid map. The BamHI-RBS-mCherry sequence effectively replaced the native gene’s protein coding sequence (shown in pink). Translational START sites are marked by the ATG codon. **C.** Promoter strength quantified by measuring cellular mCherry fluorescence in strains CJW_Bb069, CJW_Bb108 through CJW_Bb112, and CJW_Bb146. The fluorescence levels were normalized by cell area. The promoters were ranked in increasing order of the mean fluorescence values and are listed below the graph. Shown are means ± SD. Between 97 and 160 cells were analyzed per strain. EV, empty vector; A.U., arbitrary units.

We transformed these constructs into *B. burgdorferi*, imaged the resulting strains, and quantified the fluorescence level in each cell. All promoters elicited fluorescence levels above the background of the strain carrying the empty vector (Figure 3C). We noticed differences between the RNA-seq and mCherry reporter-based methods of measuring promoter strength, as detailed in the discussion. Importantly, however, the promoters we tested displayed a broad dynamic range from low (P*_0526_*) to intermediate (P*_0826_*, P*_resT_*, P*_0031_*, and P*_0026_*) to high (P*_flaB_*) strength.

### Antibiotic selection in *B. burgdorferi* using hygromycin B and blasticidin S resistance markers

Several antibiotic resistance markers have been used to perform genetic manipulations in *B. burgdorferi* and have recently been reviewed in detail (13). The most widely used today are the kanamycin (*aphI*), gentamicin (*aacC1*), streptomycin (*aadA*) and erythromycin (*ermC*) resistance genes (see Table 4) (45, 66-68). Use of several other antibiotics for selection is either ineffective (e.g., zeocin, chloramphenicol, and puromycin), discouraged due to safety concerns (e.g., tetracyclines, β-lactams, and sometimes erythromycin), redundant due to cross-resistance (several aminoglycoside antibiotics), or no longer widespread (coumermycin A_1_) due to alterations in cell physiology induced by both the antibiotic and the resistance marker (13, 67).

To expand the panel of antibiotic resistance markers that can be used in *B. burgdorferi*, we focused on two antibiotics commonly used for selection of eukaryotic cells, namely the translation inhibitors hygromycin B and blasticidin S. Rendering *B. burgdorferi* resistant to them does not pose a biosafety concern, as these antibiotics are not used to treat Lyme disease. We found that hygromycin B and blasticidin S prevented *B. burgdorferi* growth in liquid culture at concentrations of 200 and 5 μg/mL, respectively (Table 4). For resistance cassettes, we used the *E. coli* gene *hph* (also known as *aph(4)-Ia*), which encodes a hygromycin B phosphotransferase, and the *Aspergillus terreus* gene *bsd*, which encodes a blasticidin S deaminase (69-71). We codon-optimized these genes for translation in *B. burgdorferi* and placed them under the control of the strong P*_flgB_* promoter on a shuttle vector (Figure 4A). The resulting vectors, pBSV2H and pBSV2B, also carry the rifampicin resistance gene *arr-2* of *Pseudomonas aeruginosa* (72-74), which encodes a rifampicin ADP-ribosyltransferase. *B. burgdorferi* is naturally resistant to rifampicin (75, 76), but the use of rifampicin for selection in *E. coli* instead of the more expensive blasticidin S and hygromycin B antibiotics reduces the cost of generating and propagating the vectors in *E. coli*.

**Figure 4.**
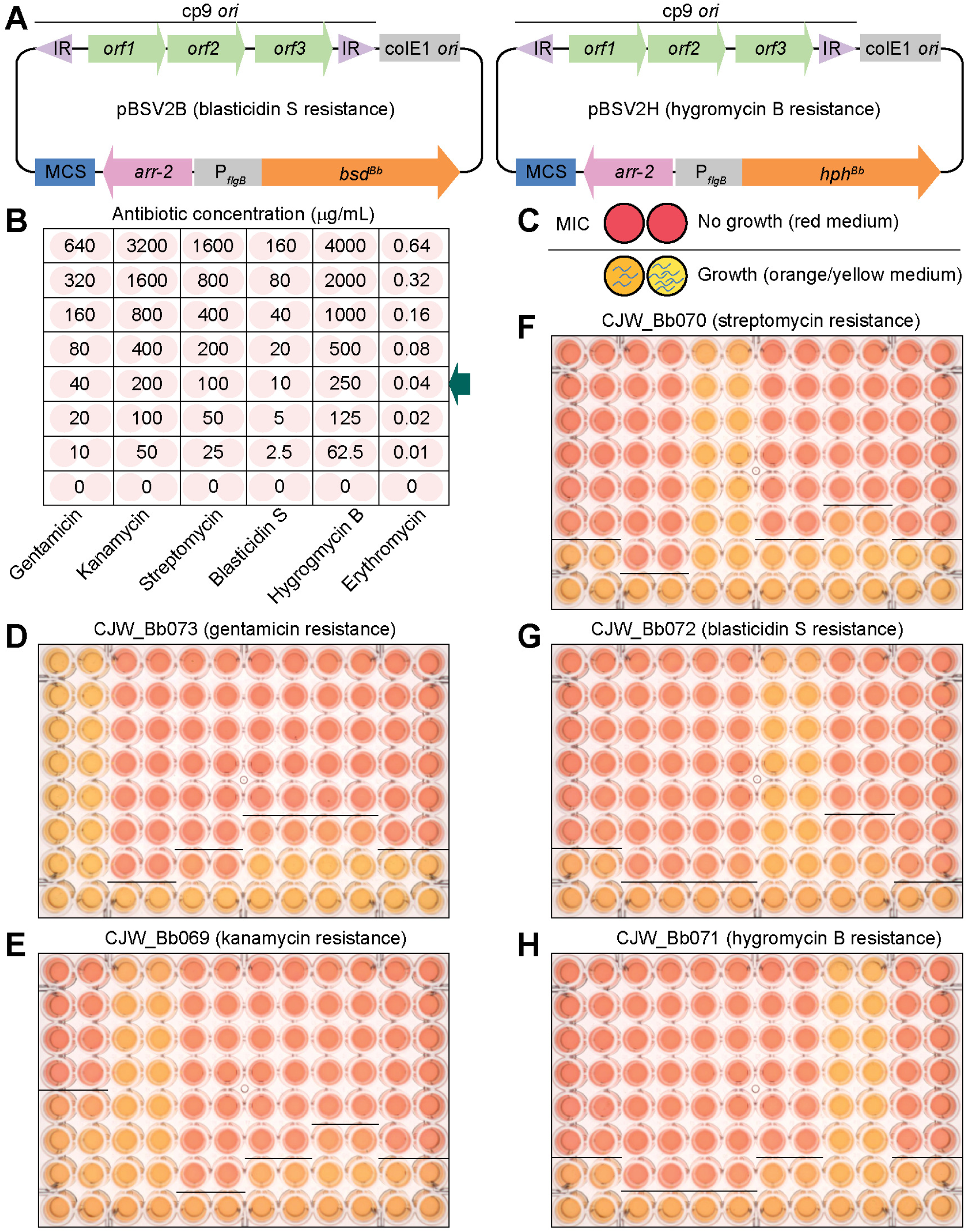
Characterization of blasticidin S and hygromycin B resistances in *B. burgdorferi*. **A.** Maps of shuttle vectors pBSV2B and pBSV2H. IR, inverted repeats; cp9 *ori*, origin or replication of *B. burgdorferi* plasmid cp9; colE1 *ori, E. coli* origin of replication; MCS, multicloning site; *arr-2*, rifampicin resistance gene for selection in *E. coli*; P*_flgB_, B. burgdorferi* flagellar rod operon promoter; *bsd^Bb^, B. burgdorferi* codon-optimized blasticidin S deaminase-encoding gene; *hph^Bb^, B. burgdorferi* codon-optimized hygromycin B phosphotransferase-encoding gene. The maps are not drawn to scale. **B.** Plate map showing the final antibiotic concentrations used for cross-resistance testing. Each concentration was tested in two adjacent wells. Concentrations routinely used for selection are indicated by the arrow. **C.** Schematic representation of color change of the growth medium from red (absence of spirochete growth) to orange/yellow (presence of spirochete growth). A line marks the boundary between growth and no growth in an antibiotic concentration series. The lowest antibiotic concentration that blocked growth was identified as the minimal inhibitory concentration (MIC). **D.-H.** Susceptibility test of each resistance-carrying strain to various antibiotic concentrations according to the plate layout shown in B. The plates were incubated to allow for growth-dependent acidification of the medium and change in phenol red pH indicator color from red to orange and yellow, as depicted in panel C. Images were obtained using colorimetric imaging of the individual plates. MIC boundaries are marked by dark lines. The strains used are listed above each image.

*B. burgdorferi* strains obtained by transforming pBSV2B or pBSV2H into B31 e2 grew readily in cultures containing 10 μg/mL blasticidin S or 250 μg/mL hygromycin B, respectively. We used these strains to test whether the antibiotic resistance cassettes encoded by these vectors conferred any cross-resistance to the often-used antibiotics kanamycin, gentamicin, streptomycin, and erythromycin. In parallel, we performed reciprocal tests using B31 e2-derived strains that carried a kanamycin, gentamicin, or streptomycin resistance cassette. Each strain was grown in the presence of two-fold serial dilutions of each antibiotic (Figure 4B). Each dilution series was centred on the concentration routinely used for selection with each of the tested antibiotics (Figure 4B, arrow). We incubated all cultures for at least four days and then inspected each well for growth by dark-field imaging. A well was considered to be growth-positive if we detected at least one motile spirochete after scanning a minimum of five fields of view. In addition, we further incubated the plates to allow for growth-dependent acidification of the medium. This pH change is easily detected as a change in the color of the medium from red, denoting no growth, to orange or yellow, denoting various degrees of growth (Figure 4C-H) (67). We confirmed that wells with the lowest antibiotic concentration at which the medium remained red also did not contain motile spirochetes. This concentration was taken to represent the minimum inhibitory concentration, or MIC (Figure 4C, black line). Whenever we exposed a strain to the antibiotic to which it carried a resistance gene, we readily detected growth at all antibiotic concentrations tested (Figure 4D-H), highlighting the efficacy of each resistance marker. Importantly, we did not detect any major cross-resistance between the five resistance markers and the six antibiotics tested (Figure 4D-H). One exception was the kanamycin-resistant strain CJW_Bb069, which was able to grow in the presence of as much as 40 μg/mL gentamicin (Figure 4E), a concentration routinely used for gentamicin selection (67). A slightly higher amount of gentamicin (80 μg/mL) was, however, sufficient to kill this kanamycin-resistant strain (Figure 4E). This low level of cross-resistance may thus necessitate use of a higher dose of gentamicin for selection if the parental strain is already kanamycin-resistant.

**Figure 5.**
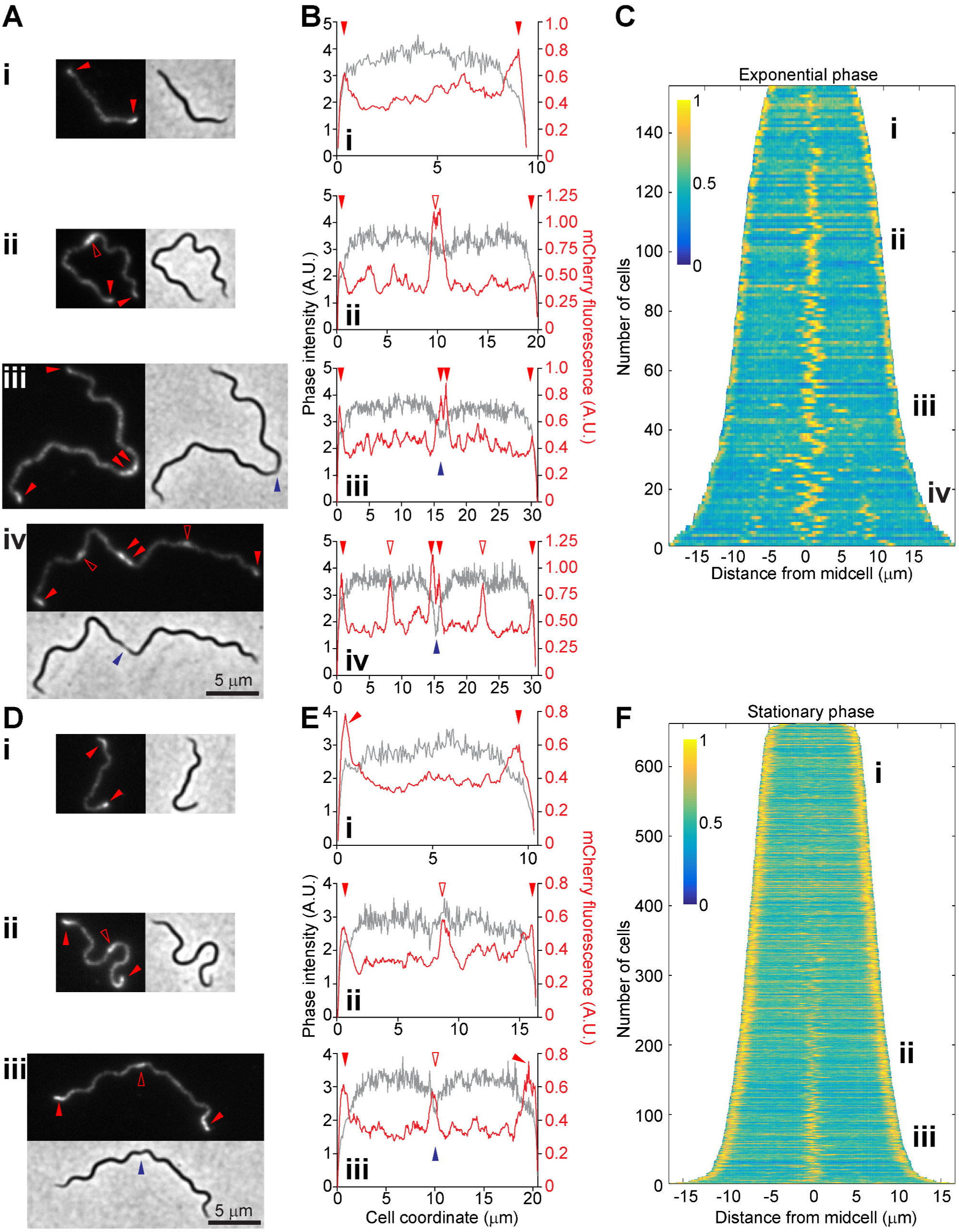
Localization of BB0323 using an mCherry fusion. **A.** Micrographs of cells of strain CJW_Bb173 imaged in exponential phase. Shown are mCherry fluorescence and phase contrast images. All images have the same magnification. Signal accumulation patterns are as follows: i, bipolar; ii, bipolar and midcell in the absence of obvious constriction in the phase contrast image; iii, bipolar and midcell in the presence of midcell constriction; and iv, bipolar, midcell in the presence of deep cell constriction, and at 1/4 and 3/4 positions along the cell length. **B.** Signal quantification along the cell length for cells shown in panel A. The mCherry signal is depicted in red, while the phase contrast signal is shown in gray. **A** and **B**. Polar localizations and midcell localizations flanking a deep constriction site are marked by filled red arrowheads. Midcell localizations are otherwise marked by empty red arrowheads. Indentation in the phase contrast signal is marked by blue arrowheads. **C.** Demograph depicting the localization of BB0323-mCherry in a population of exponentially growing cells of strain CJW_Bb173. See text for a detailed description. i-iv depict the regions on the demograph where cells with the localization patterns highlighted in panels A and B are located. **D**. Micrographs of cells of strain CJW_Bb173 imaged in stationary phase. Shown are mCherry fluorescence and phase contrast images. All images have the same magnification. Signal accumulation patterns are as follows: i, bipolar; ii, bipolar and midcell in the absence of cell constriction in the phase contrast image; and iii, bipolar and midcell in the presence of midcell constriction. **E.** Signal quantification as a function of cell length for cells shown in panel D. The mCherry signal is depicted in red, while the phase contrast signal is shown in gray. D and E. Filled and empty red arrowheads mark polar and midcell localization, respectively, whereas the blue arrowhead shows cell constriction. **F.** Demograph depicting the localization of BB0323-mCherry in a stationary phase population of cells of strain CJW_Bb173. i-iii depict the regions on the demograph where cells with the localization patterns highlighted in panels D and E are located.

### Subcellular localization of a *B. burgdorferi* LysM domain-containing protein

To highlight the usefulness of our newly generated *B. burgdorferi* molecular reagents, we fused the gene encoding mCherry to the 3’ end of *bb0323* to create a C-terminal fluorescent fusion. The resulting construct was placed under the control of the intermediate-strength promoter P*_0826_*. BB0323 is an important lipoprotein that is required for *B. burgdorferi*’s natural infection cycle through the tick and the mammalian reservoir (27). This lipoprotein is proteolytically processed into an N-terminal domain mediating outer membrane stability and cell separation, and a C-terminal fragment containing a peptidoglycan-binding LysM domain (26, 28, 77). The N- and C-terminal fragments interact with each other and were proposed to help anchor the outer membrane to the peptidoglycan (28).

We found that the BB0323-mCherry fusion displays striking localization patterns that vary predictably with the culture growth phase (Figure 5). In an exponentially growing culture, the shortest cells (likely newly born cells) displayed a patchy distribution of the BB0323-mCherry signal along the length of the cell, accompanied by accumulation of this signal at the cell poles (Figure 5A.i and 5B.i). In longer cells (i.e., later during the cell cycle), these patchy and bipolar localizations were accompanied by accumulation of the signal at midcell (Figure 5A.ii and 5B.ii). These midcell localization events coincided with future division sites (6), though signal accumulation at midcell could be detected in the absence of obvious cell constriction (Figure 5.A.ii and 5B.ii). Midcell localization persisted through cell constriction, with the fluorescent signal often becoming split into two intensity peaks that flanked the cell constriction site, as shown by the indentation in the phase contrast signal (Figure 5A.iii and 5B.iii). Upon complete cell separation, these pairs of midcell intensity peaks presumably form the polar signals of daughter cells. In a subset of deeply constricted cells, BB0323-mCherry also accumulated at the 1/4 and 3/4 locations along the cell length (Figure 5A.iv and 5B.iv), which represent the midcell positions and future division sites of the still-connected daughter cells.

The cell cycle coordination of these localization patterns was confirmed by demograph analysis of static images of an asynchronous population (Figure 5C). In this representation, each horizontal line represents the distribution of the BB0323-mCherry fluorescent signal along the length of a single cell, as depicted in a heat map. Cells are sorted by their lengths to approximate cell cycle progression.

In a stationary phase culture, the same localization patterns were observed, except for the disappearance of the signal at the 1/4 and 3/4 cell positions. We assume that the slower growth rates in stationary phase ensure that daughter cells fully separate before BB0323-mCherry begins to accumulate at midcell. Slower growth rates in stationary phase may also account for the delayed accumulation of midcell signal during this phase (Figure 5F) relative to exponentially growth (Figure 5C).

## DISCUSSION

We have undertaken this work to facilitate microscopy-based investigations of the biology of the Lyme disease agent *B. burgdorferi*. We expanded the available molecular toolkit by characterizing antibiotic resistance markers, fluorescent proteins and constitutively active promoters not previously used in this organism.

Alongside the commonly used kanamycin, gentamicin, streptomycin, and erythromycin selection markers, the addition of hygromycin B and blasticidin S resistances as useful selection markers will provide more flexibility in designing genetic modifications. A wider array of non-cross-resistant selection markers is particularly important in the absence of a streamlined method to create unmarked genetic modifications in this bacterium (13). Currently, in infectious *B. burgdorferi* strains, an antibiotic resistance marker is commonly used to inactivate the restriction modification system encoded by the *bbe02* locus on plasmid lp25. This inactivation increases the efficiency of transformation with shuttle vectors. It also helps maintain this plasmid in the cell population during *in vitro* growth through selective pressure (78-81). This latter point is essential for maintaining a strain’s infectivity, as linear plasmid lp25 is essential *in vivo*, but is often rapidly lost during genetic manipulations and growth in culture (82, 83). A second resistance marker is often used to inactivate a gene of interest, either by targeted deletion or by transposon insertion mutagenesis. A third resistance marker is needed for complementation, either at the original locus, or *in trans*. Additional markers are needed if two genes are to be inactivated and complemented simultaneously, or if several protein localization reporters need to be expressed both simultaneously and independently.

Today’s cell biology investigations often rely on microscopy studies using fluorescent protein fusions. Prior to our work, green and red fluorescent proteins have been the reporters of choice in *B. burgdorferi* microscopy studies (Table 3), and only a handful of subcellular localization and lipoprotein topology studies had been performed using these tools (13-19). We have expanded the palette of fluorescent proteins that can be used in this bacterium by adding several proteins with properties highly desirable for imaging and localization studies. These fluorescent proteins are among the brightest of their classes (20, 40, 43), and their spectral properties render them appropriate for simultaneous multi-color imaging of up to four targets. For the most part, they are also monomeric, as all of the *A. victoria* GFP, CFP, and YFP variants that we have generated carry the A206K mutation (41). Using monomeric fluorescent proteins may be important to prevent artifactual intermolecular interactions, (e.g., (41, 84, 85)). Should the weakly dimeric versions of these proteins be required for specific applications, the A206K mutation can be easily reversed by site-directed mutagenesis. Furthermore, the superfolder variants of these proteins may facilitate tagging when the folding of the fusion protein is otherwise impaired (40). In addition, unlike EGFP, which does not fold in the periplasm of diderm bacteria when exported through the Sec protein translocation system, sfGFP does fold in this compartment (86). It can therefore be an alternative to mRFP1 and mCherry for tagging periplasmic and outer surface-exposed proteins. This is particularly relevant for the study of *B. burgdorferi* since this bacterium expresses an unusually large number of lipoproteins that are localized on the cell surface or in the periplasmic space (87).

In addition, although dimeric, iRFP may serve as a useful *in vivo* marker, and may be preferable to GFP and RFP. Excitation light penetrance in live tissues is better in the farred/near-infrared region of the spectrum than in the blue-shifted regions used to excite GFP and RFP. Furthermore, tissue autofluorescence in this spectral region is lower, which further facilitates imaging (88, 89). Lastly, the levels of biliverdin found in animal tissues are in the low milimolar range, with healthy human plasma containing 0.9-6.5 μM biliverdin (90). In our hands, such biliverdin levels are sufficient to elicit maximal fluorescence of *B. burgdorferi*-expressed iRFP. Furthermore, iRFP has been successfully used to label the bacterium *Neisseria meningitidis* for *in vivo* imaging (91). Altogether, these considerations suggest that imaging in mice using iRFP-expressing *B. burgdorferi* should be feasible.

We also characterized promoters of low and intermediate strengths and demonstrated that variable degrees of constitutive gene expression can be easily achieved in *B. burgdorferi*. The relative order of promoter strength, as quantified using the mCherry reporter (Figure 3C), largely matched the order of the expression levels of the corresponding genes in culture (Figure 3A) (64), with the exceptions of P*_0526_* and P*_0826_*. While P*_0526_* had an intermediate strength as measured by RNA-seq, it was the weakest when tested using our reporter system. In contrast, P*_0826_* was the weakest promoter based on RNA-seq data, but displayed intermediate strength in our experiments. These differences may stem from strain differences or from our use of short DNA sequences of 129 to 212 bp, which presumably contain minimal promoter sequences. Any native regulatory elements located further upstream of these short promoter sequences are thus absent in our reporter plasmids. Differences in expression levels may also be caused by reporter expression from circular shuttle vectors. The native P*_0526_* and P*_0826_* sequences are located on the chromosome and differences in DNA topology, including supercoiling, between the chromosome and the circular plasmids are known to affect gene expression in *B. burgdorferi* (92, 93). Finally, it is worth noting that while both *bb0526* and *bb0826* encode leaderless transcripts, *bb0826* has a secondary transcriptional start site located 54 bp upstream of the translational start site (65). This difference may also partly explain the observed promoter strength mismatch between the native gene and reporter fusion. Regardless of the reason for these discrepancies, these promoters will facilitate complementation and localization studies where medium and low gene expression levels may be required.

To demonstrate the usefulness of these molecular reagents, we used them to generate and express an mCherry fusion to the LysM domain-containing protein BB0323. LysM domain-containing proteins, including BB0323, have been shown to bind the peptidoglycan layer (28, 94). Strikingly, BB0323-mCherry localized at the cellular poles and at future division sites at midcell throughout most of the cell cycle. Near the end of the cell cycle, before cells separate, BB0323-mCherry also often accumulated at 1/4 and 3/4 cell positions corresponding to the division sites of future daughter cells. We previously demonstrated that the midcell and the 1/4 and 3/4 positions represent regions of active peptidoglycan synthesis in *B. burgdorferi* (6). Perhaps BB0323 accumulates at sites of peptidoglycan growth because they differ in chemical composition. A more likely alternative is that these peptidoglycan regions are multi-layered. A higher local peptidoglycan concentration would provide a denser binding platform for BB0323, resulting in its accumulation. In fact, cryo-electron tomography imaging of *B. burgdorferi* has revealed multiple layers of peptidoglycan at division sites (4), as well as a thick peptidoglycan layer at the poles (95). Accumulation of BB0323 at sites of multi-layered peptidoglycan would be reminiscent of the LysM-containing protein DipM in the α-proteobacterium *Caulobacter crescentus*, which localizes at zones of multi-layered peptidoglycan, including division sites and poles, via its LysM domains (96-98). Importantly, the striking cell cycle-coordinated localization of BB0323 demonstrates that proteins with critical functions are spatially distributed in *B. burgdorferi*, highlighting a layer of regulation that has been poorly explored in spirochetes.

In summary, our study describes novel molecular tools that we hope will aid investigations in the Lyme disease field and spur further progress in the study of this medically important and highly unusual bacterium.

## DATA AVAILABILITY

Sequences of all the plasmids constructed in this study are available upon request. The DNA sequences of the various genes that were codon-optimized for expression in *B. burgdorferi* are provided in the Supplemental Material. The MATLAB code used to process cell fluorescence data is also provided as Supplemental Material.

## ACCESSION NUMBERS

DNA sequences of codon-optimized genes have been deposited at GenBank under accession numbers MH644044 through MH644053.

## SUPPLEMENTAL MATERIAL

Supplemental Material is available in the accompanying document. It contains detailed plasmid construction methods, a list of oligonucleotide primer sequences used in this study, DNA sequences of genes that were codon-optimized for translation in *B. burgdorferi*, a record of cell numbers for each figure describing quantitative fluorescence data, MATLAB code used in this study, and supplemental references.

## ACKNOWLEDGEMENT

We thank Nicholas Jannetty for help with cloning experiments and Dr. Bradley Parry for help with the computational analyses. We are also grateful to Dr. Brandon Jutras and the members of the Jacobs-Wagner laboratory for helpful discussions and/or critical reading of the manuscript.

